# Lipids as currency in ecological interactions: Competition and facilitation between two lipid scavenging parasitoids

**DOI:** 10.1101/2020.03.11.987453

**Authors:** Mark Lammers, Tim A. M. van Gorkum, Stefanie Hoeijmans, Ken Kraaijeveld, Jeffrey A. Harvey, Jacintha Ellers

## Abstract

Interspecific interactions in nature often revolve around the acquisition of nutrients. Depending on the organisms’ metabolic requirements, competition for specific essential nutrients may occur, which selects for increased abilities to monopolize, consume and store these nutrients. Lipid scavengers are organisms that rely on exogenous lipid acquisition as they lack the ability to synthesize fatty acids *de novo* or in sufficient quantity. Most parasitoid insects are lipid scavengers: they obtain all required lipids by feeding on their hosts as larvae. Here we study the nutritional ecology of competitive interactions between native *Nasonia vitripennis* and introduced *Tachinaephagus zealandicus*. While the former was already known to lack lipogenesis, we show that *T. zealandicus* also relies on host lipids. The interactions between the two species were studied using competition experiments, in which oviposition of *T. zealandicus* on a host was followed by multiparasitism by *N. vitripennis*. The outcome of competition was determined by the duration of the time lag between oviposition events. *N. vitripennis* was superior when arriving 3 days after oviposition by *T. zealandicus*. In contrast, 9 days after oviposition of *T. zealandicus* we observed complete reversal, and no *N. vitripennis* offspring were able to develop. Only when *N. vitripennis* laid eggs 15 days after *T. zealandicus* oviposition, both species could emerge from the same host. However, *N. vitripennis* realizes only 10% of its potential fitness at this time point because prior parasitization by the gregarious *T. zealandicus* compartmentalizes the host resources, limiting the spread of *N. vitripennis’* venom. This study shows that successful reproduction of *N. vitripennis* at 15 days was achieved by hyperparasitizing, a capability that provides a fitness benefit to *N. vitripennis*, as it extends the time window that hosts are available for parasitization. Choice tests with hosts at different time intervals after *T. zealandicus* oviposition revealed a partial mismatch in *N. vitripennis* females between competition avoidance and offspring performance, which may be linked to the limited co-evolutionary time between native and introduced species. We discuss our results in the context of nutritional ecology and, specifically, the role of lipids in ecological interactions.

## Introduction

Interspecific interactions in nature often revolve around the acquisition of nutrients, for example through facilitation or competition among interacting species (Stuart Chapin et al. 2012). Competitive interactions can occur if multiple organisms compete for access to the same nutrients from the same source (Gause 1934; Tilman 1982). Depending on the organisms’ metabolic requirements, competition may involve monopolization of different types of nutrients. Well-known examples include strategies whereby plants compete to obtain nitrogen-containing nutrients in nitrogen-limited environments (Moreau et al. 2019), and shifts in nutrient limitations among herbivore life-stages affecting foraging strategies (Richard and de Roos 2018). Competitive pressure depends on the balance between the availability of nutrients as well as the metabolic requirements of the competing organisms, with the strongest competitive interaction expected when the nutrients that are competed for are limited but essential for metabolic functions. Essential nutrients are metabolites that the organisms cannot synthesize themselves but are required for normal physiological function (Burr et al. 1932; Mazumdar and Striepen 2007); hence these nutrients have to be taken up from an external source. Species differ in their exogenous requirement for nutrients such as vitamins, amino acids and lipids, which are essential for some species but not for others (Mazumdar and Striepen 2007). Competitive interactions between species for essential nutrients are likely to impose selection on their ability to monopolize, consume and store these nutrients.

An increasing number of species has been shown to rely on exogenous lipid acquisition as they lack the ability to synthesize fatty acids *de novo* or in sufficient quantity (Ellers et al. 2012; Keymer and Gutjahr 2018). Across several kingdoms of life lipid scavenging or lipid parasitism has evolved, which involves the transfer of lipids from host in a symbiotic interaction, for instance mycorrhizal fungi (Luginbuehl et al. 2017; Keymer and Gutjahr 2018), parasitic fungi (Xu et al. 2007), parasitic protists (Schwelm et al. 2015), apicomplexan parasites (Mazumdar and Striepen 2007), and the bacterium *Spiroplasma* (Herren et al. 2014) are all known to lack lipogenic abilities. However, the highest frequency of lipid scavenging is found among parasitic insects: parasitoid wasps, flies and beetles (Giron and Casas 2003; Visser and Ellers 2008; Visser et al. 2010). These parasitoids obtain all required lipids by feeding on their hosts as larvae, which likely relaxed selection on autotrophic fatty acid production (Lahti et al. 2009), eventually leading to full dependency on host lipids (Ellers et al. 2012).

Parasitoids develop in or on a host, killing it in the process (Godfray 1994). The developing parasitoid larva is thus dependent on the resources available from a single host to successfully complete development (Vinson and Iwantsch 1980; Sequeira and Mackauer 1992; Harvey 2005). Field studies have occasionally found up to ten or even more parasitoid species per host species (Price 1972; Elzinga et al. 2007; Oishi and Sato 2008; Kostenko et al. 2015; Abell et al. 2016; Šigut et al. 2018), so competition between parasitoid species attacking the same host is expected to be common in nature (Price 1972; Slansky 1986). Acquisition of sufficient lipid reserves during the larval period is crucial to parasitoid fitness as most (but not all) parasitoid species lack lipogenic abilities (Visser et al. 2010), and lipids of freshly emerged adults are necessary for egg maturation and survival (Ellers 1996). However, whether and how competitive interactions have shaped the lipid acquisition strategy of parasitoid species is currently unknown.

Here we study the nutritional ecology of competitive interactions between parasitoid wasps attacking the same host, focusing on how differences in host utilization strategy between the species affect their competitive abilities. The jewel wasp *Nasonia vitripennis* is an ectoparasitoid that lays its eggs on blowfly host pupae in carrion, where it may regularly encounter hosts that have previously been parasitized by earlier-arriving larval-pupal parasitoid species from the genera *Tachinaephagus, Alysia* and *Aphaereta*, some of which are capable of parasitizing up to 20% of available hosts in open semi-natural field sites (Voss et al. 2009), whereas up to 48% of hosts can be parasitized in urban biotopes (Frederickx et al. 2013). *Tachinaephagus zealandicus* (Hymenoptera: Chalcidoidea) is an endoparasitoid, ovipositing multiple eggs in the larvae of a blowfly host on carrion (Olton and Legner 1974). These parasitoid species nowadays coexist on at least four continents (Carvalho et al. 2004; Turchetto and Vanin 2004; Voss et al. 2009; Frederickx et al. 2013) It is therefore likely that *N. vitripennis* regularly encounters hosts previously parasitized by *T. zealandicus* and may have evolved adaptations to dominate in intrinsic competition for host resources (Price 1972) or to avoid competition altogether by discriminating against parasitized (Cusumano et al. 2016). Consequently, they are therefore likely to compete for host resources under certain circumstances (van Velzen et al. 2016).

When encountering a parasitized host pupa, *N. vitripennis* can either multiparasitize the host and compete with the first parasitoid for nutrients (Harvey et al. 2013), or hyperparasitize the parasitoid larvae already growing in the hosts, thus consuming nutrients that have been metabolized by the first parasitoid (Sullivan and Völkl 1999; Harvey et al. 2009) or exploit both simultaneously (Harvey et al. 2011). It is known that *N. vitripennis* is able to hyperparasitize larval parasitoids, e.g. this is occasionally observed for the solitary *Alysia manducator* (Graham-Smith 1919; Altson 1920; Peters and Abraham 2010). However, whether this species can at least partially hyperparasitize facultatively on the gregarious *T. zealandicus* is unknown. Hyperparasitism would confer higher fitness if *T. zealandicus* would be capable of fatty acid synthesis or would be more efficient in acquiring lipids from the host. Lack of lipogenesis has been confirmed in *N. vitripennis* (Rivero and West 2002; Visser et al. 2012; but see Prager et al. 2019), but it is hitherto unknown whether *T. zealandicus* possesses any lipogenic capabilities. Furthermore, the duration of the time lag between arrivals of either species (i.e. the developmental stage of *T. zealandicus*) is likely to have an effect on the reproductive success and lipid acquisition of *N. vitripennis* (Harvey et al. 2011; Zhu et al. 2016).

In the present study we investigate (1) whether *T. zealandicus* shows lack of lipogenesis, (2) whether the developmental stage of *T. zealandicus* has an effect on *N. vitripennis’* offspring emergence success and fitness, (3) whether *N. vitripennis* is able to hyperparasitize on *T. zealandicus*, (4) whether *N. vitripennis* can detect the presence of *T. zealandicus* larvae inside a potential host, (5) if females prefer to oviposit on a host that has not been previously parasitized by *T. zealandicus*, and (6) whether there is a match between offspring lipid acquisition and host preference (Jaenike 1978).

**Figure 1.**
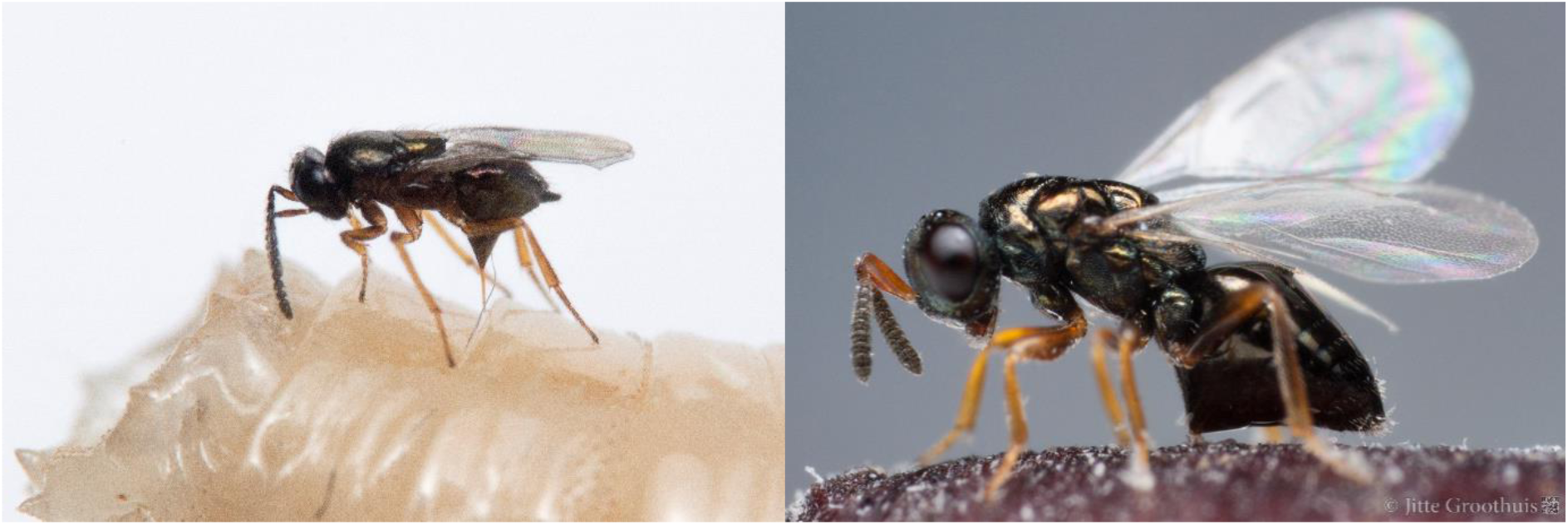
Images of adult females of the two parasitoid species studied here: *Tachinaephagus zealandicus* ovipositing on a host larva (left) and *Nasonia vitripennis* ovipositing on a host pupa (right). Pictures by Jitte Groothuis.

## Methods

### Study organisms

*Nasonia vitripennis* (Hymenoptera: Chalcidoidea: Pteromalidae) is a cosmopolitan idiobiont ectoparasitoid (Whiting 1967) and a lipid scavenger (Visser et al. 2012). The females oviposit on fly puparia, which she finds on carcasses and in birds’ nests (Whiting 1967; Peters 2010). After the eggs hatch, the first instar larvae will move away from the egg shell and begin to feed on the host by puncturing the host skin with their mandibles. The larvae imbibe the body fluid of the host and will remain in the same position, unless disturbed, until it is fully grown (Whiting 1967). Development from egg to adult takes about 22 days at 20°C. In our experiments, we used the isofemale strain AsymCX, which had been reared on *Calliphora vomitoria* at the Vrije Universiteit Amsterdam for 52 generations prior the experiments.

*Tachinaephagus zealandicus* (Hymenoptera: Chalcidoidea: Encyrtidae) is a koinobiont endoparasitoid with unknown lipogenic abilities, originating from Australasia (Olton and Legner 1974; Subba Rao 1978). It was introduced to some areas as a biological control against carrion flies, spread independently to other locations, and is now found globally (Olton and Legner 1974; Ables 1977; Ferreira de Almeida et al. 2002; Geden and Skovgård 2014; Peters 2014). The eggs are oviposited in the fly larvae, but the parasitoid development only starts after the host has pupated (Olton and Legner 1974). The strain “HHx” of *T. zealandicus* used in this study was reared from *Parasarcophaga caerulescens* larvae, gathered on 30 August 2014 from Oostvoorne, the Netherlands (H. Huijbregts, pers. comm.). This strain was reared in the lab for 5 generations prior to the experiments. The development from egg to adult takes about 32 days at 20°C.

### Testing lipogenic abilities of *T*. *zealandicus*

Freshly emerged wasps were randomly allocated to one of two treatments: ‘Emergence’, in which wasps were killed by freezing at −20°C on the day of emergence, or ‘Fed1wk’, which had *ad libitum* access to a 20% (w/v) sucrose solution for one week. We measured the lipid content of wasps from the Emergence treatment, and wasps that survived the Fed1wk treatment. We followed a modified protocol of David et al. (1975; Ellers 1996). Wasps were first individually checked for body integrity under a microscope (Leica WILD M8). The wasps were each placed in a labelled glass vial (Lenz Laborglas, Wertheim, Germany), freeze-dried for 2 days and subsequently weighed on a microbalance (Mettler Toledo UMT2, d=0.1 µg). Next, 4 mL di-ethyl ether was added to each vial in order to dissolve all lipids from the wasp’s body. After 2 days, the wasps were removed from the ether and dipped in fresh ether to wash off any residue. All wasps were freeze-dried again and weighed on the same microbalance. The lipid content of each wasp was calculated as the dry weight before ether extraction minus the dry weight after ether extraction. Afterwards, all wasps were checked for body integrity. Wasps that missed any body parts were removed from the analysis because this would bias the calculation of lipid content.

Differences in lipid content between treatments were analyzed using ANCOVA with treatment as independent variable and fat-free dry weight as a measure for body size as covariate. An increase in lipid content after one week of feeding is evidence for lipogenic abilities, while lack of lipogenesis would be inferred if the lipid content was highest at emergence.

### Design and timing of the treatments in competition experiments and choice tests

Competitive performance of either species was tested in competition experiments and host preference was assessed with choice tests. To obtain consistently parasitized hosts for each test in sufficient quantity, we prepared an excess of tubes with one *T. zealandicus* female, one *C. vomitoria* larva and some saw dust, at 20 °C. At the same time, an equal number of *C. vomitoria* larvae were added to a glass rearing jar with a layer of sawdust to allow them to pupate; these were the non-parasitized hosts used as controls in the experiments.

*T. zealandicus* parasitized hosts were offered at various stages of development to *N. vitripennis* females. Host pupae in the ‘Early’ treatment contained egg or first instar larva of *T. zealandicus*, in the ‘Middle’ treatment they contained second to third instar larva of *T. zealandicus*, and in the ‘Late’ treatment the host was completely consumed by *T. zealandicus* larvae, just before these started to pupate.

To determine the exact timing of Middle and Late treatments after host pupation, we first measured the time required for *T. zealandicus* to reach the different developmental stages at 20°C. 100 females were allowed to oviposit individually on single hosts. 25 hosts were dissected every five days after oviposition and developmental stage of the parasitoids inside was recorded. Based on the resulting developmental curves (see results), the timing of the Early, Middle and Late treatments were determined at 3, 9 and 15 days after *T. zealandicus* parasitized the host, respectively. These treatment timings were implemented in both the competition experiments and the choice tests.

*N. vitripennis* females that are inexperienced in laying eggs will take a longer time to find and parasitize a host than experienced females (Rivers and Denlinger 1994). Therefore, females were ‘trained’ prior to the experiments following standard protocols by giving them an oviposition experience. Briefly, fresh females were placed in a plastic tube with 20-25 fly pupae with demi water and honey offered on the plug, and left at 20°C for one day. At least three hours prior to the start of the experiment, the hosts were removed from the tube to give the females time to recuperate before the start of the experiment, as *N. vitripennis* females need 1-4 hours after oviposition to recover, before doing so again (Edwards 1954; King and Rafai 1970). This experience was given to all *N. vitripennis* females prior to both the competition experiments and choice tests.

All competition experiments and behavioral observations were verified by dissections of parasitized hosts. All experiments were performed at 20°C, 75% relative humidity and a 16:8 L:D light regime.

### Competition experiments

To determine the competitive strength of each species at each time point, 120 pupated hosts (in two blocks of 60, separated by one day) parasitized by *T. zealandicus* were divided over the three competition treatments described above and put separately in a 75×23.5mm plastic tube with a styrofoam plug. At the appropriate timing for the treatment a fed and experienced *N. vitripennis* female of 3 to 4 days old was added. The females were allowed to oviposit for 24 hours at 20°C, after which the parasitized hosts were kept at 20°C. Four control treatments were performed in parallel in order to disentangle the effects of host age and multiparasitism on the performance of *N. vitripennis* and *T. zealandicus*, each with the same replication as in the competition treatments. Three control treatments (Control_Early_, Control_Middle_ and Control_Late_) had non-parasitized hosts of ages matched to the respective competition treatments. To each host a single *N. vitripennis* female was added which was allowed to parasitize for 24 hours. Control_Late_ was found to give an unexpected, but trivial, zero-fitness result for *N. vitripennis*, as the host flies already emerged from the pupae two days before the wasp was supposed to oviposit. To test the performance of *T. zealandicus* when parasitizing alone, a control treatment (Control_Tz_) was performed in which each host was parasitized by one *T. zealandicus* female as above, but without later addition of *N. vitripennis*.

We measured several fitness components of emerging offspring: emergence success, brood size, development time, and offspring lipid content. The number of successfully emerging wasps of either species was scored for each host in all treatments. Development time was measured in days between oviposition and emergence of the first individual of each species. The lipid content of one random female per emerging brood of *N. vitripennis* was measured using the methods described above. The total brood mass of *T. zealandicus* was measured because the brood size varied notably: brood sizes ranging from 2 to 102 offspring were found. Total brood mass was measured by freeze-drying the broods in Eppendorf tubes for 48 hours, and their dry weight was measured including the tubes. After the initial weighing, the wasps were removed from the tubes, which were then cleaned with a soft brush and weighed again. The dry weight of the brood was obtained be subtracting this latter weight from the initial measurement.

No *N. vitripennis* offspring emerged in the Middle treatment. In order to confirm that *N. vitripennis* actually oviposited on the hosts offered in the competition experiments of this treatment, an extra experiment was performed similar to the Middle treatment. The host was made only partially available to the ovipositing female by putting the *T. zealandicus*-parasitized host in a foam plug with a pupa-sized hole in it, so that only the posterior end of the pupa was exposed. This made it easier to locate *N. vitripennis* eggs when opening the cocoon on the posterior end.

In a separate experiment we compared the effects of *N. vitripennis* venom injection against mechanical damage alone, on survival of *T. zealandicus* larvae. 40 hosts parasitized by *T. zealandicus* in the Late developmental stage were split over two treatments: either they were offered individually to a *N. vitripennis* female as above in a plug with a pupa-sized hole in it for a period of 24 hours, after which the female was removed; or we applied a control treatment where we inflicted only mechanical damage by puncturing the host between the second and third segment from the posterior end with a sterilized microneedle with diameter 32 - 126 µm (measured from tip to thickest point), slightly bigger than *N. vitripennis’* ovipositor which is approximately 24 µm thick. 48 hours after the start of the experiment all cocoons were carefully opened at the exposed area. Any *N. vitripennis* eggs were removed from the sting site to exclude effects induced by the developing *N. vitripennis* offspring. The hosts with *T. zealandicus* larvae were placed gently inside a transparent gelatin capsule (size 1, SVM Grondstoffen, De Meern, the Netherlands), to protect them from injury and desiccation. They were allowed to develop to adulthood, after which all *T. zealandicus* were killed by freezing. Not all wasps complete development and emerge successfully. The host carcasses were dissected and the numbers of developed and undeveloped *T. zealandicus* were counted.

### Choice tests

We performed a series of choice test to determine whether *N. vitripennis* females discriminate between *T. zealandicus* parasitized hosts and non-parasitized hosts at each of the three treatment time points. These choice tests were conducted by placing an experienced *N. vitripennis* female and two fly pupae in a 55mm Petri dish without vents. The pupae were placed 2 cm apart and at equidistance from the center of the dish in a tiny drop of water to prevent them from rolling around. In the Early (n=60), Middle (n=60), and Late (n=30) treatments, one of the hosts was parasitized by *T. zealandicus*, and the other was not parasitized. The non-parasitized hosts for the Late treatment were 11 days old instead of 15 days, because *C. vomitoria* emerge from the pupae after 13 days. The parasitized host was placed to the left or to the right of the center of the Petri dish at random for each separate trial. As a control treatment, females were given access to two non-parasitized hosts (n=40).

We scored for each trial 1) whether the wasp landed on any host; 2) on which host the wasp landed first; 3) what behaviors the wasp performed on that first host (Edwards 1954); 4) whether the wasp chose to oviposit on any host; and 5) which host was finally chosen for oviposition. Once a female had completed the full behavioral sequence, it was considered to have made a choice and then she was removed from the Petri dish. Wasps that had not performed this full behavioral sequence after 1.5 hours were considered not to have made a choice. Afterwards all hosts were stored individually in labelled Eppendorf tubes and stoppered with a foam plug.

To confirm that the hosts that should have been parasitized by *T. zealandicus* indeed contained parasitoid larvae, the hosts were dissected and investigated under a microscope (Leica WILD M8). If larvae were visible inside the host, or if the host was disintegrated (the effect of *T. zealandicus* venom) it was marked as parasitized. If the host was not parasitized by *T. zealandicus*, the choice the individual had made was invalid and the data from this trial excluded from the analysis. In addition, this dissection served to verify oviposition by *N. vitripennis* on the host. All other hosts were kept in a climate chamber at 25°C and checked for emergence of *N. vitripennis*, as emerging offspring are direct evidence of successful oviposition. If no wasps had emerged from the pupa after the expected time for emergence of *N. vitripennis*, the pupa was dissected. In all these cases the host was dead and dried out. As there was no way to tell if the host had been alive at the time of the experiment, the data observed from these replicates were removed from the analysis. Based on these criteria we removed the data from 11, 9 and 1 choice trials from the treatments Early, Middle and Late, respectively.

### Data analyses and statistics

All analyses were performed in R version 3.2.1 (R Development Core Team 2015).

In the competition experiments, emergence of any number of individuals of a species was scored as a success for that species, i.e. both species can be successful on the same host (host-sharing). Emergence success of *N. vitripennis* and *T. zealandicus* was tested for significant differences using an overdispersion-corrected binomial Generalized Linear Model with treatment as independent variable for both species separately.

To determine the effects of competition on the fitness of the emerged offspring of *N. vitripennis* and *T. zealandicus*, we tested for differences in their fitness components. For *N. vitripennis* we tested for significant differences in the fitness components brood size, development time and lipid content. Differences in developmental time were tested using a Kaplan-Meier survival analysis with a Log-rank test to test for differences between the fitted curves. Differences in brood sizes were compared using a non-parametric Kruskal-Wallis test due to the non-normality of the data. As a posthoc test we compared each of the treatments pairwise with a Wilcoxon rank sum test with Holm-Bonferroni p-value adjustment (R Development Core Team 2015). The lipid content of wasps were compared using a Generalized Linear Mixed Model (GLMM, nlme package) with lipid content as dependent variable and fat-free body weight as co-variable, treatment as independent variable, and block as random variable. A post-hoc Tukey-test was done to compare the differences between each of the treatments when the GLMM showed a significant result.

As a final analysis of the costs and benefits of competition we calculated the estimated fitness for each competition treatment expressed as the average amount of lipids acquired per brood in that treatment. Lipid acquisition was calculated by multiplying mean brood size with mean per capita lipid content. Variances were summed accordingly. Wasps have optimal fitness on recently-pupated hosts without competitor (Whiting 1967), i.e. the Control_Early_ treatment is expected to provide the highest lipid content. 95% confidence intervals of the amount of lipid acquired were calculated for each treatment. Estimates of lipids acquired with non-overlapping confidence intervals were considered to be significantly different.

For *T. zealandicus* we compared the total dry weight of the brood as a measure of fitness. Differences in the total dry weight of the successfully emerged offspring were tested using a GLMM with log-transformed weights (to meet assumptions of normality) as dependent variable, treatment as independent variable, and block as random variable. A posthoc Tukey-test was performed to compare the differences between each of the treatments when the GLMM showed a significant result. Furthermore, we compared the numbers of undeveloped larvae from the separate experiment where broods were attacked by *N. vitripennis* or mock-injected with a microneedle using a one-way ANOVA.

The choice tests were analysed in four subsequent steps. (1) The number of trials where a wasp did not land on any host were tested for significant differences between treatments using an overdispersion-corrected binomial Generalized Linear Model with treatment as independent variable. These trials were excluded from subsequent data analyses. The same statistical test was applied for the number of trials where a wasp did not make a choice for any host, i.e. the wasp landed on at least one host, but did not oviposit. If the GLM was found to give significant results, a posthoc test using Dunnett contrasts was applied. (2) The number of trials where wasps chose to land on the parasitized host was tested for each treatment separately with a Binomial test against a null expectation of random choice, i.e. the probability of choosing the parasitized host was set at 0.5. (3) For each of the behaviors we tested whether the odds ratio of performing the next behavior was less than one using Fisher’s Exact Test separately for each combination of treatment and host first landed on. If the odds ratio to proceed with the next behavior is significantly less than one, this is indicative of rejection of the host after performing a certain behavior. (4) The number of trials where wasps chose to oviposit on the non-parasitized host was tested for each treatment separately with a Binomial test against a null expectation of random choice, i.e. the probability of choosing the parasitized host was set at 0.5.

## Results

### No increase in *Tachinaephagus zealandicus* lipid content after one week of feeding

The lipid content of *T. zealandicus* females (Figure 2) that fed *ad libitum* on a sucrose solution for one week was significantly lower than in freshly emerged wasps (ANCOVA, F_1,20_=31.12, p<0.001) and increased with body size (p<0.001).

**Figure 2.**
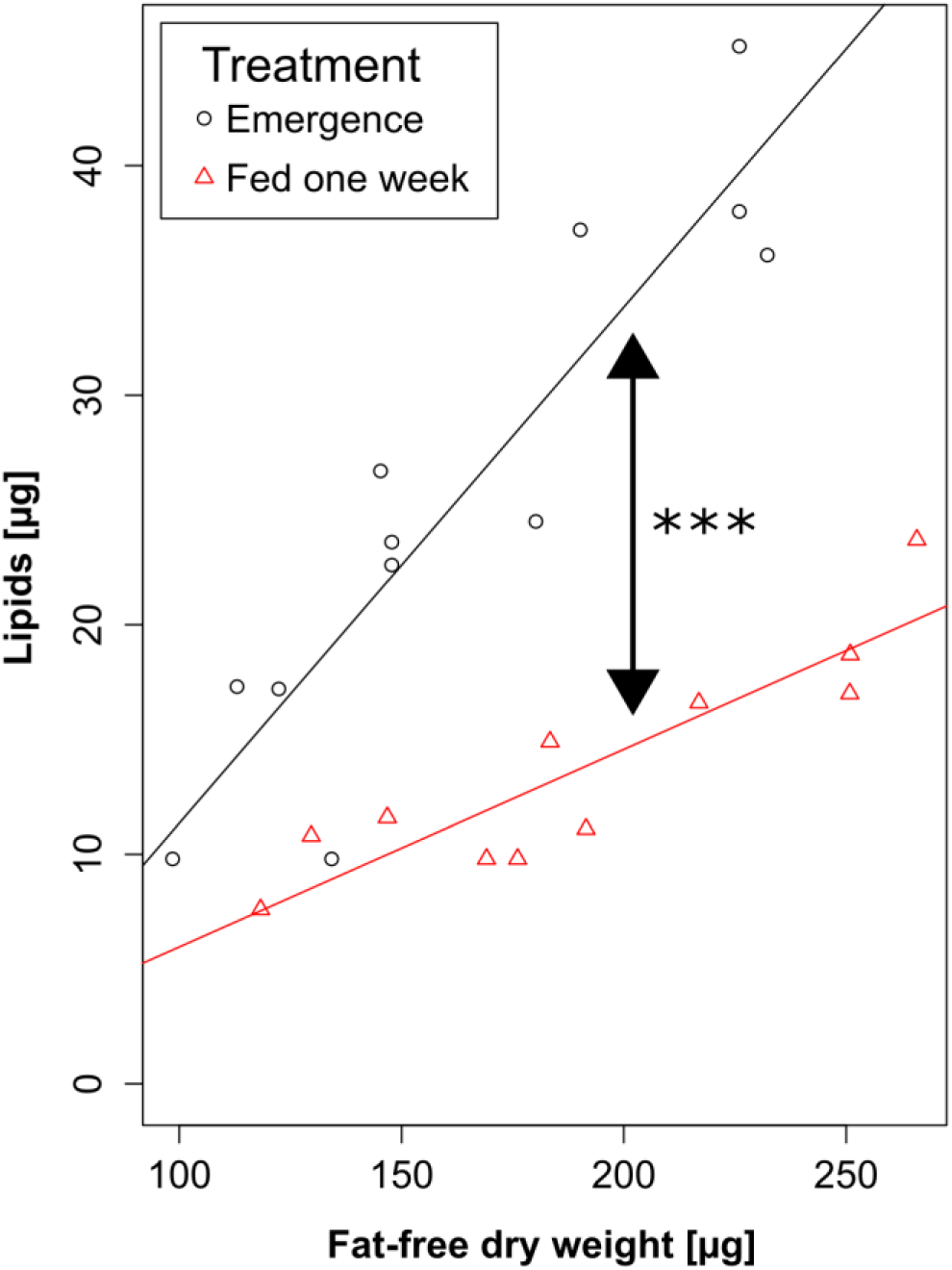
*T. zealandicus* is a lipid scavenger: Lipid content of *T. zealandicus* was highest at emergence and had significantly lower after one week of feeding on a sucrose solution.

### *Tachinaephagus zealandicus* development and timing of treatments

All hosts pupated within 3 days after oviposition by *T. zealandicus* on the host larvae. The wasp’s developmental curves from egg, through the larval instars and pupal stage, to adult are plotted in figure 3A. Figure 3B shows the derived timing of the treatments for the competition experiments and choice tests.

**Figure 3.**
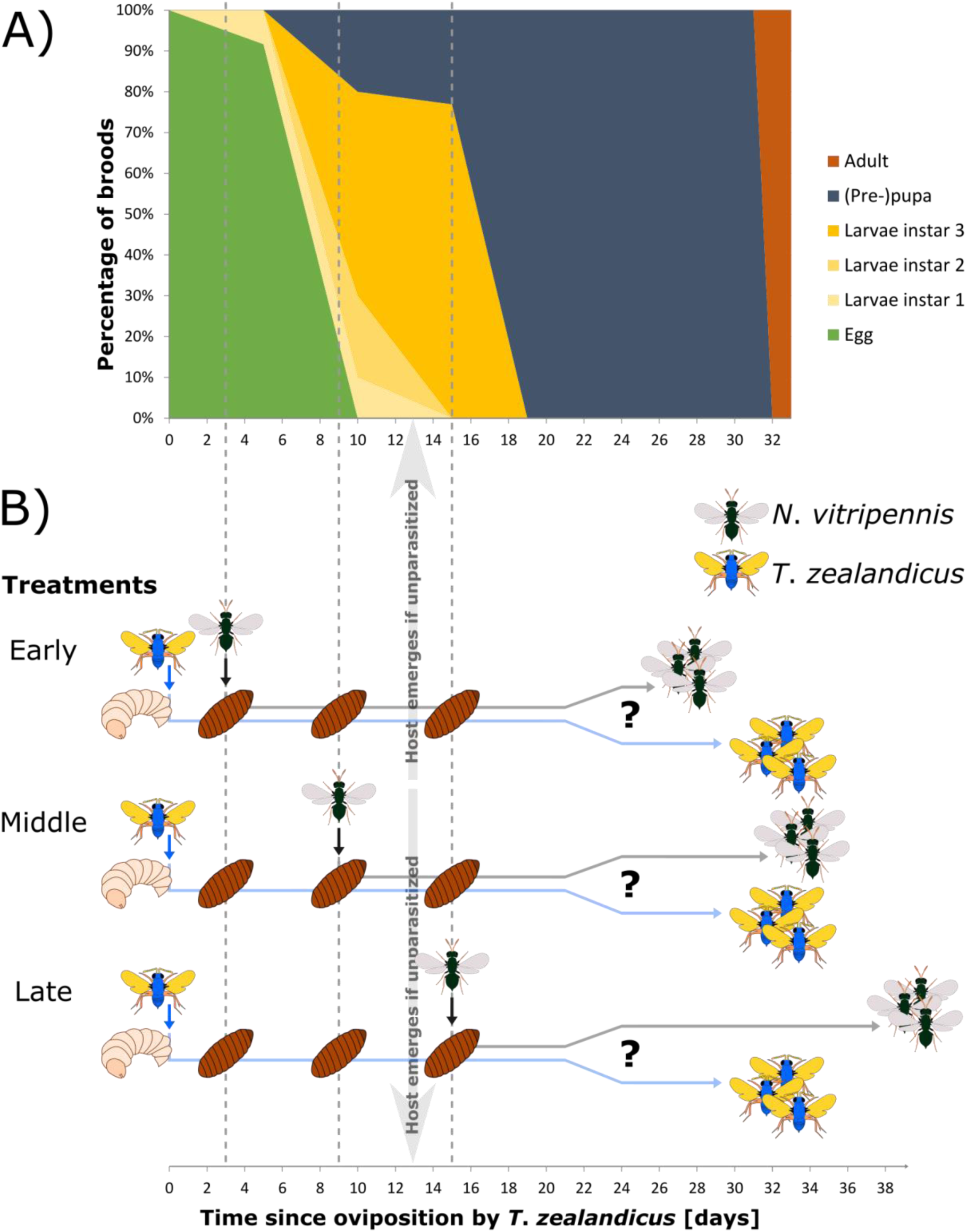
Development of *T. zealandicus* at 20°C through all life stages, and the timing of treatments for competition and choice tests. A) The development of *T. zealandicus* from oviposition to adult emergence. 25 parasitized hosts were dissected on day 5, 10, 15 and 19 to determine the percentage of broods per developmental stage. Vertical dotted lines at 3, 9 and 15 days represent the timing of the Early, Middle and Late treatment, respectively. B) The experimental setup where *N. vitripennis* is introduced to host parasitized by *T. zealandicus* during three different developmental stages of *T. zealandicus* inside the host. The Early treatment shows *N. vitripennis* introduced just after the parasitized host has pupated, the Middle treatment shows the introduction of *N. vitripennis* during a later larval instar phase of *T. zealandicus*, and finally, the Late treatment shows *N. vitripennis* introduction after the host has been fully consumed by *T. zealandicus*. The approximate moment where unparasitized hosts emerged is indicated. Control treatments (i.e. either species’ success without competition) are not depicted.

### Emergence success of both species

The emergence success of both species in the different treatments indicates which of the two species is dominant in a specific treatment (Figure 4). *N. vitripennis* emerged from 65% of hosts in the Control_Early_ to 70% of hosts in the Control_Middle_. The presence of *T. zealandicus* in the Early treatment had no effect on the emergence success of *N. vitripennis* compared to the success in Control_Early_ (Binomial test, df=39, p=0.869, 95% confidence interval for the probability of success = [0.509, 0.814]). Not a single *N. vitripennis* emerged in the Middle treatment, in sharp contrast to the Control_Middle_ (Binomial test, df=39, p<0.0001, 95% confidence interval for the probability of success = [0.000, 0.088]). In the Late treatment, *N. vitripennis* emerged from 10% of the hosts, significantly more than in the controls (see methods; Binomial test, df=39, p<0.0001, 95% confidence interval for the probability of success = [0.028, 0.236]).

**Figure 4:**
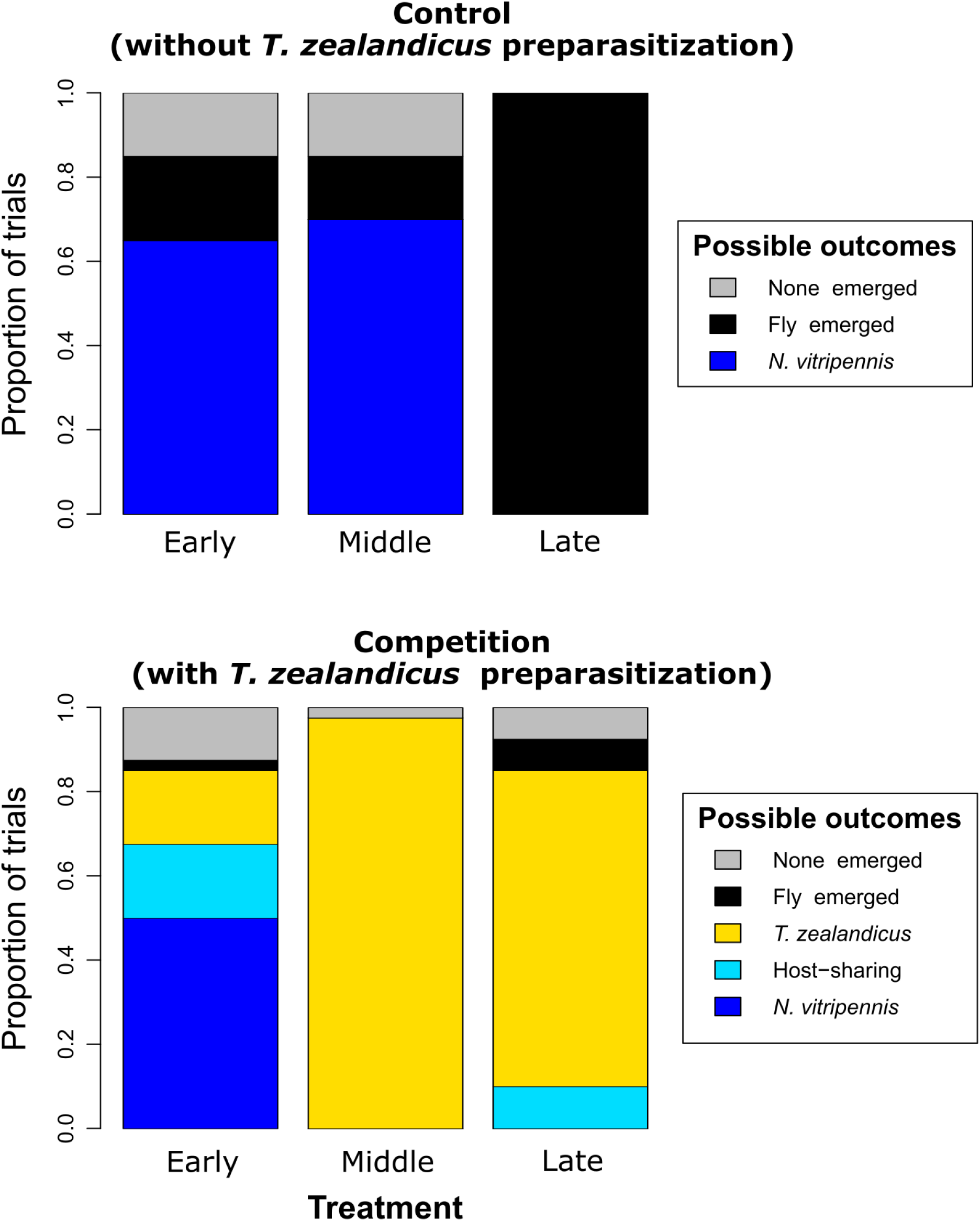
Emergence success of both species of parasitoids per treatment. The proportion of trials where either of the two parasitoid wasp species successfully emerged from the different competition treatments and controls is shown on the y-axis (n=40). Possible outcomes of the trials are: only *N. vitripennis* emerged (dark blue), only *T. zealandicus* emerged (yellow), both parasitoid species emerged (light blue), the host fly emerged (black) or none emerged (grey) from the host pupae. See the main text for a description of significant differences

*T. zealandicus* emerges from 90% of hosts in Control_Tz_. Multiparasitism by *N. vitripennis* significantly reduced its success to 35% in the Early treatment (Binomial test, df=39, p<0.0001, 95% confidence interval for the probability of success = [0.206, 0.517]). However, emergence success of *T. zealandicus* was similar to Control_Tz_ in the Middle (Binomial test, df=39, p=0.180, 95% confidence interval for the probability of success = [0.868, 0.999]) and Late treatments (Binomial test, df=39, p=0.287, 95% confidence interval for the probability of success = [0.702, 0.943]).

Of the 57 hosts from which *N. vitripennis* successfully emerged, 7 also produced *T. zealandicus*. Four of these were in the Late treatment, i.e. they produced both parasitoid species, while none produced *N. vitripennis* only. No host resources are available anymore in the Late treatment, hence in these cases *N. vitripennis* can only have hyperparasitized on *T. zealandicus*

Note that the proportion of hosts where nothing emerged were all similar to the *Tachinaephagus*-parasitized control after correcting for multiple comparisons (Binomial tests, df=39, p>α/6). See figure 4 for an overview of the emergence success of both parasitoid species.

### Effects of competition on fitness components of *N*. *vitripennis*

Since no offspring of *N. vitripennis* emerged in the Middle treatment, and that wasps did not get an oviposition opportunity in the Control_Late_, there is no quantitative comparison possible for the fitness components in these two treatments. Eggs were found on 15 out of 24 *T. zealandicus*-parasitized hosts in a separate experiment, hence oviposition by *N. vitripennis* on parasitized hosts was confirmed for the Middle treatment.

The egg-to-adult development time of *N. vitripennis* (Figure 5A) differed between treatments (ANOVA, F_3, 81_=10.108, p<0.0001). Development was significantly longer in the Control_Middle_ than in Control_Early_ (Tukey contrasts, p=0.002), and even longer development in the Late treatment than in the other treatments (Tukey contrasts, p=0.018).

**Figure 5:**
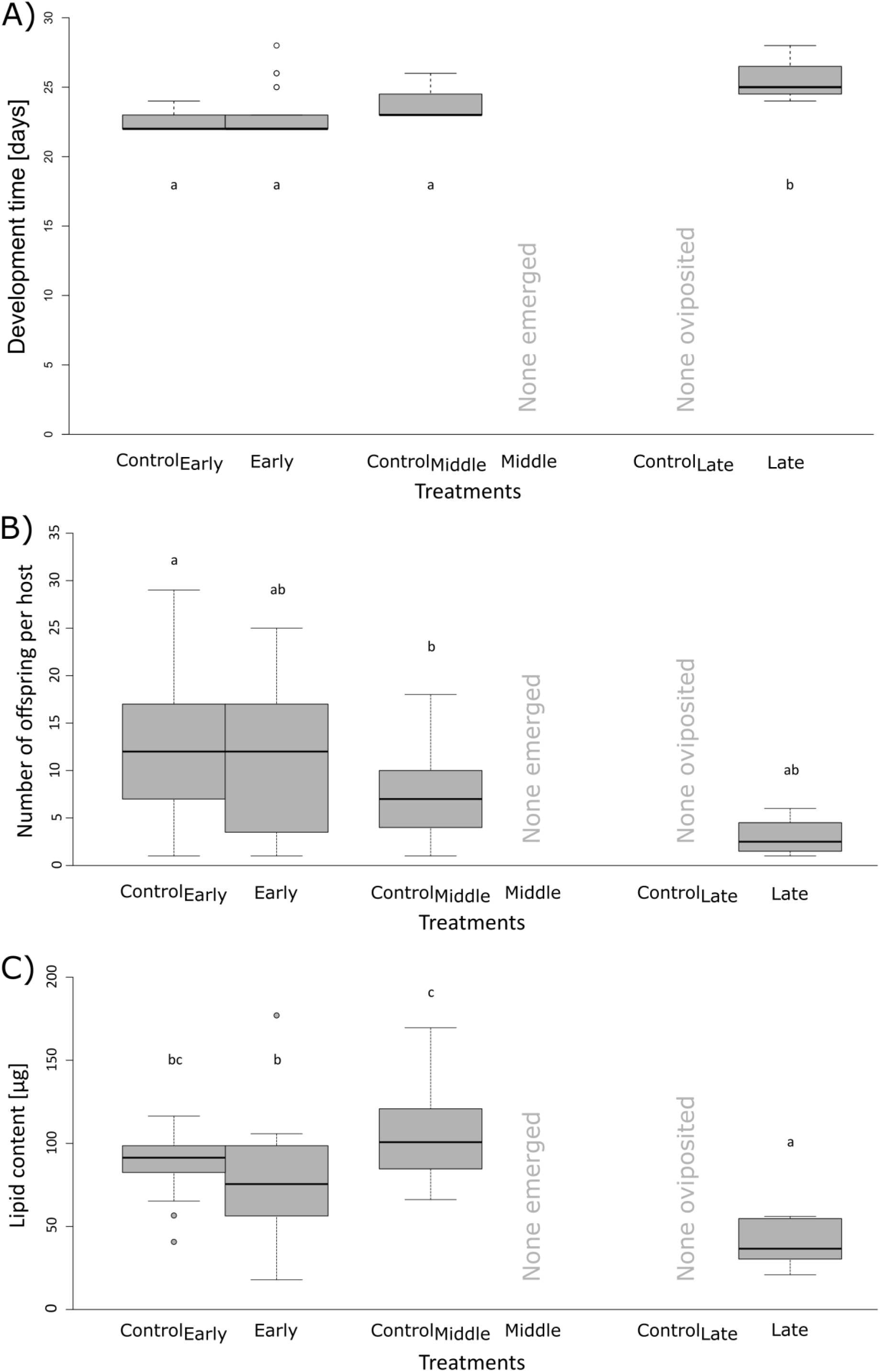
Fitness effects of competition on fitness components of N. vitripennis. A) Egg-to-adult development time of *N. vitripennis* per treatment. B) Brood size of *N. vitripennis* brood sizes per treatment. C) Lipid content of individual *N. vitripennis* offspring per treatment. In all plots, lower case letters denote significant differences.

Brood size of *N. vitripennis* (Figure 5B) was significantly different between the treatments (Kruskal-Wallis test, χ2=11.81, df=3, p=0.008). Post-hoc tests revealed a significantly larger brood size in Control_Early_ than in Control_Middle_ (Pairwise Wilcoxon rank sum test, p=0.042), suggesting that recently pupated hosts are potentially the best stadium for *N. vitripennis* brood size. The lowest brood size (1-6 offspring per host) is found in the Late treatment, a difference which is approaching significance when compared to the Control_Early_ treatment (Pairwise Wilcoxon rank sum test, p=0.061). There was no significant difference between the brood sizes of the Early treatment relative to all other treatments.

The lipid content of *N. vitripennis* females (Figure 5C) differed significantly between treatments (GLMM, χ2=24.796, df=3, p<0.0001), with wasps in the Late treatment having significantly lower lipid content than in the other treatments (Tukey contrasts, all p<0.05). The lipid content in Control_Middle_ was significantly higher compared to the lipid content in the Early treatment (Tukey contrast, z=-2.944, p=0.017). There was no significant difference between the two control treatments (Tukey contrasts, all p>0.05).

*N. vitripennis* was not affected by competition when arriving 3 days after oviposition by *T. zealandicus*, as there was no significant difference between the Early treatment and Control_Early_ for any of the fitness components (development time, brood size, lipid content; respective Tukey contrasts, all p>0.05).

Only in the Early treatment did we observe hosts from which *N. vitripennis* as well as *T. zealandicus* emerged (i.e. host-sharing) and hosts from which either species emerged alone. Hence, we can compare the fitness of *N. vitripennis* between shared hosts and non-shared hosts for the Early treatment. All fitness components were lower in wasps from shared hosts than in wasps from non-shared hosts: development time was longer (GLMM, χ2=6.991, df=1, p=0.008), brood size was smaller (GLMM, χ2=11.690, df=1, p<0.001), and their lipid content was lower (GLMM, χ2=4.99, df=1, p=0.025).

We estimated the total lipids (in µg) acquired per treatment from a given host for *N. vitripennis* (Figure 6A). The 95% confidence intervals for lipid acquisition in Control_Early_, Control_Middle_ and the Early treatment were mostly overlapping: [717.9, 1441.4], [511.0, 1114.4], and [553.6, 1359.5], respectively. In the Late treatment the 95% confidence interval for lipid acquisition was significantly lower at [33.8, 204.5]. In this treatment it is 15 days after *T. zealandicus* oviposited, when the host is completely consumed and compartmentalized (see figure 6B).

**Figure 6.**
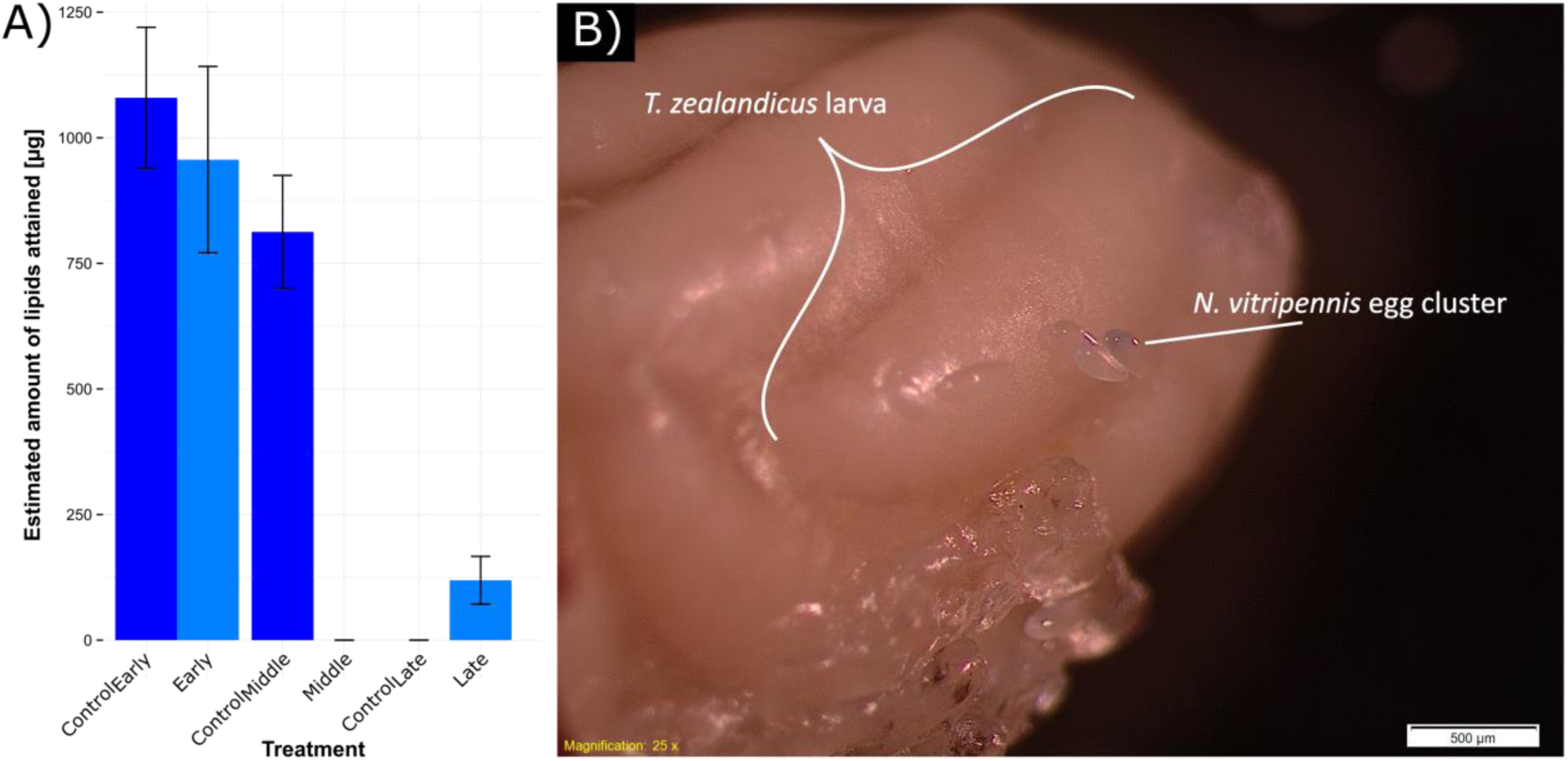
A) Estimated total amount of lipids acquired (mean±SE) per brood of *N. vitripennis* per treatment, calculated from results presented in figure 5. B) Eggs of *N. vitripennis* on fully-grown *T. zealandicus* larvae can only succeed by hyperparasitizing. Here the host cocoon is removed and 15 day old *T. zealandicus* larvae inside the host’s skin are visible (i.e. the Late treatment time point). A cluster of *N. vitripennis* eggs was found to be oviposited onto them. The host is already fully consumed by the *T. zealandicus* larvae, effectively compartmentalizing the host’s available resources.

### Effect of multiparasitism by *N*. *vitripennis* on *T*. *zealandicus*

Figure 7A shows the dry weight of *T. zealandicus* broods in each of the treatments. There was a difference among the treatments (GLMM, χ^2^=39.384, df=3, p<0.001), with the total brood mass in the Early treatment being lower than the other treatments (Tukey contrasts, all p<0.001). There was no significant difference between any of the other treatments (all p>0.05).

**Figure 7.**
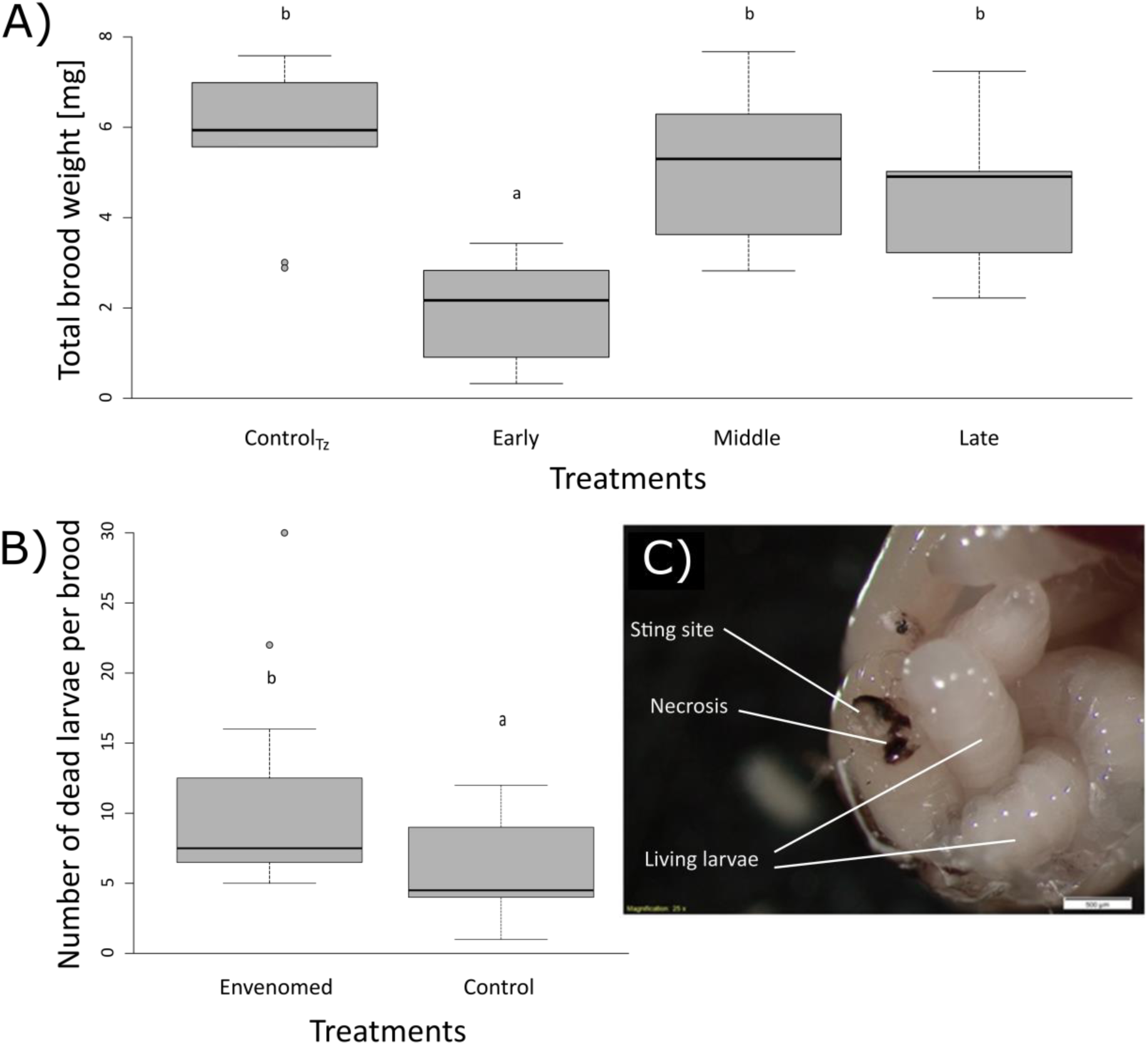
*T. zealandicus* is negatively affected by competition with *N. vitripennis*. A) Boxplot showingthe differences in total brood dry mass for *T. zealandicus* between the treatments. Lower case letters *zealandicus* larvae that were found in hosts envenomed by *N. vitripennis* and mock-stung hosts. Lower denote significant differences. B) Boxplot showing the number of development-arrested *T*. case letters denote significant differences. C) Picture of the *N. vitripennis* sting site on *T. zealandicus*larvae. Effect of the venom is visible as necrosis (black tissue) on one of the larvae. Note the lack ofeffect on other larvae in the brood.

The effect of *N. vitripennis* venom on the larvae of *T. zealandicus* caused a significantly higher number of undeveloped (dead) larvae in the envenomed treatment compared to the microneedle-injected control treatment (Figure 7B, ANOVA, F_1, 36_=7.855, p=0.008). However, many larvae survived the *N. vitripennis* venom injection and could continue their development. Figure 7C shows a photograph of typical effects of envenomation by *N. vitripennis* on *T. zealandicus* larvae.

### Host preference

The number of trials in the treatments and control where *N. vitripennis* females did not land on either host ranged from 5.0 to 14.3%, which was not significantly different among treatments (GLM, X^2^=3.030, df=3, p=0.387). While wasps showed no preference for either host to land on first in the Middle or Late treatments (Binomial tests, p=0.442 and p=0.666, respectively), 66.7% of wasps in the Early treatment preferred to land first on a host previously parasitized by *T. zealandicus* (Binomial test, p=0.044).

After this first landing, wasps can proceed with drumming, drilling and finally oviposition. We tested for each treatment if this behavioral sequence was broken off and at which behavioral step (Fig 8). The odds ratio to proceed with the next behavior was significantly less than unity in the Early treatment, as both on the *T. zealandicus*-parasitized host and on the non-parasitized host fewer wasps oviposited on the host after performing the drumming behavior (Fisher’s Exact tests, p=0.020 and p=0.002, respectively). In the Middle treatment, fewer wasps proceeded with drilling after drumming on the parasitized host (Fisher’s Exact test, p=0.025), and in the Late treatment, fewer wasps oviposited subsequent to drilling into the parasitized host (Fisher’s Exact test, p<0.0001).

**Figure 8.**
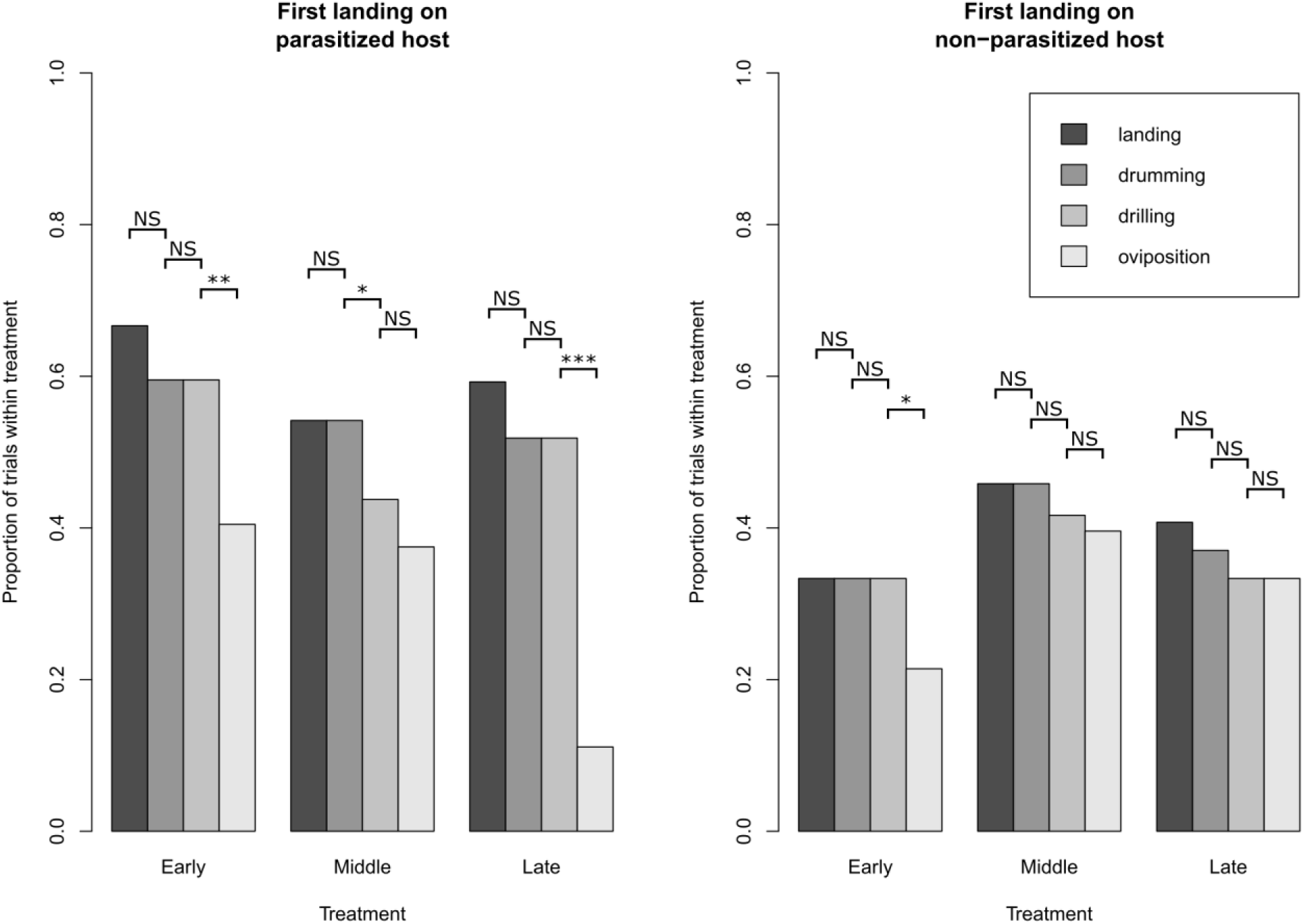
Proportions of female *N. vitripennis* that performed behaviors on the first host they landed on, split into landing first on the parasitized (left panel) versus the non-parasitized host (right panel), for each of the treatments.

After landing on at least one host, there were significant differences between treatments in the number of wasps that did not oviposit at all (GLM, X^2^=12.453, df=3, p=0.006). In the Early treatment, a significantly higher percentage of wasps did not oviposit (42.9%) compared to the control (7.9%) (Dunnett’s contrasts, p=0.029). The final choice of host for oviposition (see figure 9) did not differ from random choice in the Early or Middle treatments (Binomial tests, p=0.185 and p=0.880, respectively), while in the Late treatment there was a significant preference to oviposit on non-parasitized hosts (Binomial test, p=0.002, 95% confidence interval=[0.626, 0.953]).

**Figure 9.**
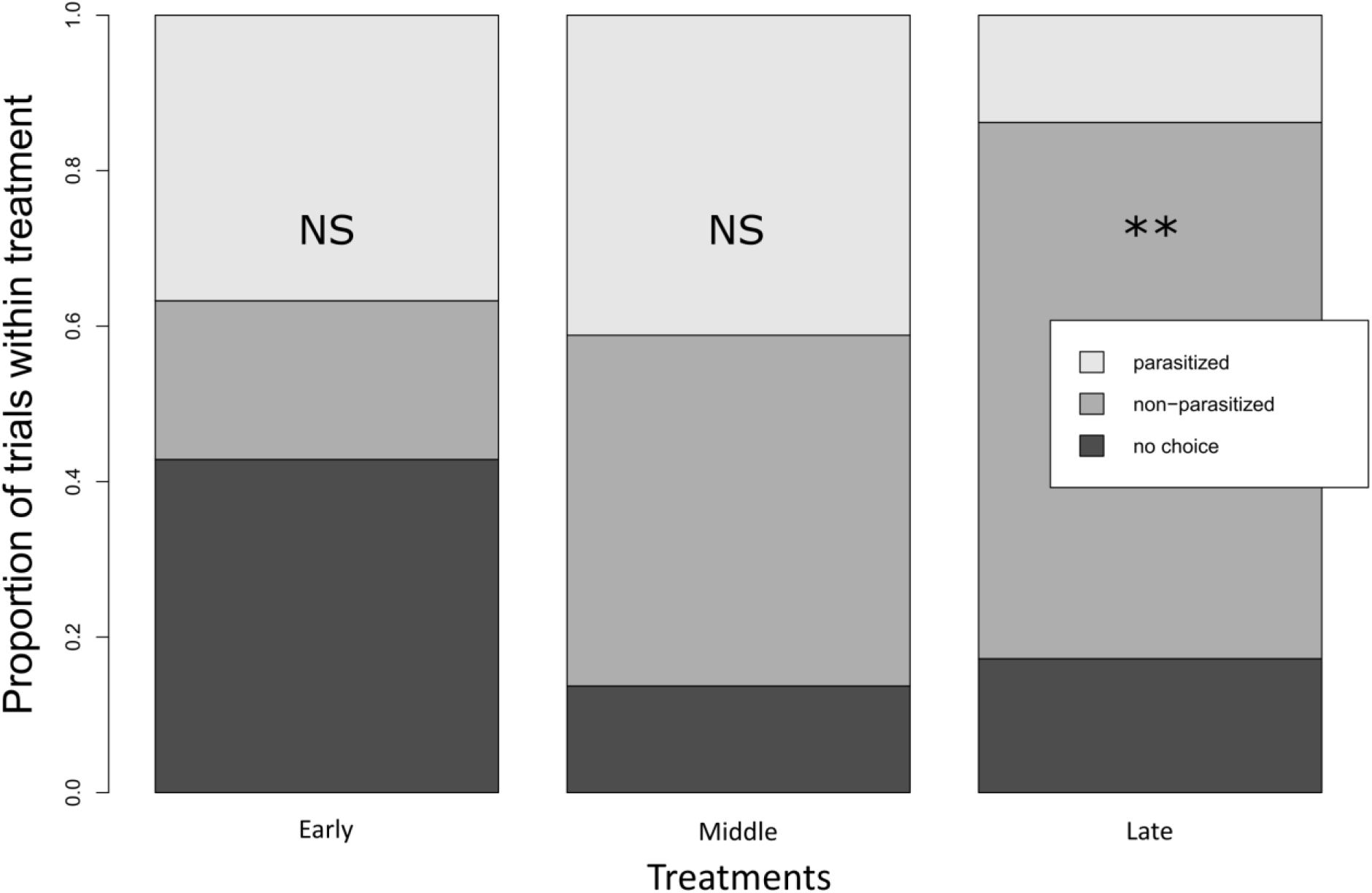
Final choice of female *N. vitripennis* after examining at least one host in each of the treatments.

## Discussion

In the present study we investigated competitive interactions for host resources between immature stages of two parasitoid wasp species. Since both species are unable to build lipid reserves from *de novo* synthesis, we expected competition over the available resources to be most intense for host lipids. As predicted, the amount of lipids acquired by the offspring of pupal parasitoid *N. vitripennis* was substantially reduced when the earlier-arriving parasitoid *T. zealandicus* had been present in the host longer. At the stage when *T. zealandicus* has fully consumed the hosts (the Late treatment), *N. vitripennis* oviposited more often in non-parasitized hosts, but at the other stages no preference was observed. Coevolution between the two species would be expected to match maternal host preference to offspring performance (Jaenike 1978). This was found in the Early treatment, where there was no preference and lipid acquisition was equal to controls. However, a match between preference and performance was not found in the Middle treatment, where wasps obtained no lipids from previously parasitized hosts, but nevertheless showed no host-discrimination. Furthermore, most wasps in the Late treatment chose to oviposit on nearly-emerging non-parasitized hosts, while the parasitized host provided them with a small amount of lipids. Below, we further discuss temporal and nutritional interaction between both parasitoids, focusing on the role lipids.

### Competitive dominance depends on time of arrival

Sequential multiparasitism of a host by the two species is a form of direct resource scramble competition that affects reproductive success of both competitors. Several studies have shown that ectoparasitoids often win the competition with endoparasitoids, especially when the latter are in the early stages of development at the time of multiparasitism (Flanders 1971; Briggs 1993; Borer 2002). In line with previous studies, *N. vitripennis* outcompeted *T. zealandicus* in most of the broods in the Early treatment, hence the emergence success of *T. zealandicus* was strongly reduced compared to controls without *N. vitripennis* oviposition. In fact, the proportion of hosts from which *T. zealandicus* emerged as the sole species was roughly equal to the proportion of hosts that was not successfully parasitized by *N. vitripennis* in the control treatment (i.e. the proportion of host from which flies emerged). This suggests that at this stage of development *T. zealandicus* mostly profits from the lack of successful parasitism by *N. vitripennis*, albeit with strongly reduced brood weight. Scramble competition between parasitoids has been reported to lead to reduced individual mean body size (Slansky 1986; Harvey 2000; Duval et al. 2018), but no significant effects of competition on the size of emerging *N. vitripennis* were found in this treatment: the total amount of lipids acquired were similar between treatments and controls where the parasitoids did and did not experience competition. However, a significant negative effect for all fitness components was found in those few broods where both species emerged from the same host (i.e. host-sharing), which indicates that scramble competition did occur in some of the broods.

*T. zealandicus* was the only species to emerge in the Middle treatment, where *N. vitripennis* oviposited on hosts containing 10 day old *T. zealandicus* larvae at which time some host resources were still unconsumed. It is unclear what kind of interaction causes complete exclusion of *N. vitripennis*. At least 60% of *N. vitripennis* females were observed to have oviposited on the parasitized hosts in a separate verification experiment (data not shown). Borer (2002) found that the increased developmental stage of the first parasitoid had a negative effect on the success of the later-arriving parasitoid when both were parasitizing the same host. Furthermore, it is possible that *T. zealandicus* larvae at this stage of its development are less sensitive to the venom of *N. vitripennis*. The successful emergence of *T. zealandicus* suggests that the majority of their larvae are not lethally affected by the venom.

*N. vitripennis* only emerged from four hosts in the Late treatment, the stage when the 15 day old *T. zealandicus* already consumed the entire host. In all these cases, they emerged from hosts that also produced *T. zealandicus* offspring. The most plausible explanation is that *N. vitripennis* is capable of facultative hyperparasitation on these gregarious *T. zealandicus* larvae, as this is the sole possible resource for the larvae at this stage. The ability to hyperparasitize is known to be present in *N. vitripennis*: it was previously found to be a facultative hyperparasitoid on the solitary endoparasitoid *Alysia manducator* (Graham-Smith 1919; Altson 1920; Peters and Abraham 2010). A study by Harvey et al. (2011) showed a similar facultative shift in trophic level by *Gelis agilis*, a solitary secondary hyperparasitoid, when it encountered *Lysibia nana* in the host *Cotesia glomerata*. In this system, multiparasitism occurred if both species parasitized within 24 hours after each other. However, *G. agilis* switched to hyperparasitization when arriving 72 hours later than *L. nana*, facultatively increasing its trophic level. Here we find for two gregarious species that at an early stage multiparasitism occurs, and later *N. vitripennis* switches to facultatively hyperparasitizing on *T. zealandicus*.

### Compartmentalization of the host by *T*. *zealandicus* prevents spread of *N*. *vitripennis* venom

Why is *N. vitripennis* not able to utilize all of the resources contained in the *T. zealandicus* larvae by hyperparasitism? Prior to ovipositing eggs on any host, the *N. vitripennis* female will inject venom into the host which causes developmental arrest and results in a series of changes in the intermediary metabolism that could benefit the *N. vitripennis* development (Rivers and Denlinger 1995; Martinson et al. 2014; Mrinalini et al. 2015). After envenomation, the eggs are laid on the surface of the integument. The gregarious *T. zealandicus* effectively divide the hosts’ resources into multiple (two to more than a hundred) compartments by consuming the entire host (Figure 6B). After removal of the host cocoon, the compartmentalization of the host by *T. zealandicus* larvae was observed to inhibit the diffusion of the venom throughout all available resources (Figure 7C). The envenomation site would therefore be the only potential site where the *N. vitripennis* larvae could obtain resources from *T. zealandicus*. These limitations of hyperparasitizing on *T. zealandicus* have consequences for *N. vitripennis’* fitness, as observed by the lower number of broods from which *N. vitripennis* emerged, the smaller brood size, the extended developmental time, and the offspring’s reduced lipid levels.

Since *T. zealandicus* is shown here to have a lipid scavenging strategy like most other parasitoids (Visser et al. 2010), hyperparasitism will not benefit *N. vitripennis* in obtaining extra lipids, as the competitor does not produce any. If anything, the conversion cost into first *T. zealandicus*’ tissue and then consumption by *N. vitripennis* is expected to reduce the available quantity of lipids. However, conversion efficiencies by hyperparasitoids of *Cotesia* are found to be surprisingly high (Harvey et al. 2006, 2009, 2015, 2016). Moreover, both species are probably selected to catabolize lipids sparingly in the adult stage, as they are both lipid scavengers. However, *N. vitripennis* emerging in the Late treatment had only half the lipid content of wasps in any other treatment.

### Mutual interference and facilitation in one interspecific interaction

In line with predictions, competition affected both parasitoid species negatively. This mutual interference was found in all treatments and especially in cases of host-sharing. Particularly in the Late treatment fitness of *N. vitripennis* was much lower. This may be due to a switch to hyperparasitizing on *T. zealandicus*, because conversion losses of resources accumulate with every additional trophic level. These apparent negative effects might seem like a deterrent for *N. vitripennis* to hyperparasitize on *T. zealandicus*. However, host-sharing also enhances host parasitism opportunities for *N. vitripennis* because the competitor’s larvae provide an extension to the time window of host availability for *N. vitripennis. Calliphora vomitoria* hosts normally complete development and emerge on days 13-14 of the experiment, whereas hosts parasitized by *T. zealandicus* extend development to at least 15 days and possibly longer. In our experiment *N. vitripennis* females in the Late treatment were able to oviposit eggs on these old hosts parasitized by *T. zealandicus*, and they successfully emerged, albeit in low numbers. In summary, the presence of *T. zealandicus* facilitates *N. vitripennis* under these specific circumstances, allowing a longer window of host availability for hyperparasitization. Although successful hyperparasitization occurred only rarely in our experiments, it is highly beneficial when no other hosts are available: A bad host is better than none at all. In the wild, *N. vitripennis* might adapt to the presence of the introduced *T. zealandicus* by improved abilities to hyperparasitize.

### A mismatch between host preference and offspring performance

One of the aims of this study was to investigate whether *N. vitripennis* prefers unparasitized over hosts previously parasitized by *T. zealandicus*, in order to avoid competition for limited (host) resources. *N. vitripennis* appears to be unable to use external cues for measuring host quality (King and Rafai 1970; Rivers et al. 2012), but probing host tissues by ovipositor drilling allows *N. vitripennis* to discriminate between dead and healthy hosts (Wylie 1958), between non-parasitized hosts and hosts parasitized by other *Nasonia* species (Ivens et al. 2009), and between hosts parasitized by conspecifics and other pupal parasitoids (Wylie 1971; Rivers 1996). Our results show that *N. vitripennis* does not discriminate between parasitized and non-parasitized hosts in the Early and Middle treatments. Only in the Late treatment did the females oviposit on the non-parasitized host significantly more often than expected by random choice. This suggests that *N. vitripennis* can only detect the *T. zealandicus* larvae when they are starting to pupate and the host tissue has been completely consumed. In the Early treatment there was a marginally significant preference to land first on the parasitized host. In the other treatments the first landing was random. This suggests that *N. vitripennis* is not capable to determine the host quality from a distance. However, after drilling into the parasitized hosts a higher proportion of *N. vitripennis* moved to the other host in the Early and Late treatment. Crucially, this is also the time that wasps form associative memories between characteristics of the environment and oviposition rewards (Hoedjes et al. 2014). In the Early treatment, first landing was biased to parasitized hosts, but final choice for oviposition was 50:50. And although first landing was random in the Late treatment, more wasps oviposited on non-parasitized hosts (note that these hosts were about to emerge as flies).

Host preference and offspring performance are predicted to co-evolve in order to maximize offspring fitness (Jaenike 1978; Cusumano et al. 2016). However, in the case of interaction with a nonnative species there may not have been sufficient time to optimize the behaviors. *Tachinaephagus zealandicus* was introduced from Australia into Denmark in 1970 as a (unsuccessful) biocontrol agent against house flies (Geden and Skovgård 2014), while *N. vitripennis* is native in Europe. This leads to an important question: Is 45 years of co-evolutionary interaction enough time for selection to optimize behavioral responses to an invader? The host flies are already attacked by a range of other parasitoid species of the genera *Aphaereta, Alysia, Muscidifurax, Spalangia, Trichopria*, and others (Frederickx et al. 2013; Mitroiu 2013; van Achterberg et al. 2020). Considering the presence of so many different species attacking the same fly hosts, it is likely that *N. vitripennis* evolved adaptations to cope with interspecific competition. These may be coopted as exaptations in its interaction with *T. zealandicus*. In our study, we found a mismatch between *N. vitripennis* host preference and offspring performance in two out of three scenarios, which suggests that these species have not had sufficient time of co-evolutionary interaction to optimize behaviors. Depending on the frequency with which interactions between species occur in nature, *N. vitripennis* might evolve to become more effective in hyperparasitizing gregarious endoparasitoids in a highly competitive environment, for example in areas with high *T. zealandicus* densities. This experiment is currently being carried out in nature, as both species spread globally and have already been found co-occurring at the same sites on four different continents (Gold and Dahlsten 1981; Bishop et al. 1996; Oliva 2008; Frederickx et al. 2013)

### Measuring lipids acquisition in ecological interactions of lipid scavengers

In this study we unravel a competitive interaction between two species of parasitoid wasps. In addition to successful emergence of offspring, we also studied lipid acquisition as the main currency of the interaction. We propose to look beyond survival and body mass as measures of fitness in future studies. Specifically, we suggest to focus on nutrients that are limiting in the examined ecological interaction (Richard and de Roos 2018). Here were presented an example of two species of parasitoid insects that are lipid scavengers and are thereby intrinsically limited in lipids. By quantifying fitness as lipids acquired we found differences between treatments that may otherwise have been missed, for example in treatments where brood sizes of *N. vitripennis* were similar, or where two fitness proxies indicated opposite effects. Measurements on the flow of specific nutrients are expected to be helpful in every system where prior knowledge is available on such nutrient limitation (e.g. Wilder et al. 2013; Herren et al. 2014; Paredes et al. 2016; Keymer and Gutjahr 2018).

## Acknowledgements

Natalie Wagner provided a helping hand during a crucial point of the competition experiments. Robin van der Slikke performed several of the choice tests as part of a high school assignment. Jitte Groothuis is thanked for the photography of both parasitoid wasp species. This work was supported by a grant from The Netherlands Organization for Scientific Research [NWO, VICI grant number 865.12.003].

